# Single-Cell Metabolic Profiling in a Glioblastoma Co-culture Model Using AP-MALDI-based Mass Spectrometry Imaging

**DOI:** 10.64898/2026.01.14.699449

**Authors:** Une Kontrimaite, Kei F. Carver Wong, Sandra Martínez-Jarquín, Phoebe McCrorie, Ruman Rahman, Dong-Hyun Kim

## Abstract

Mass spectrometry imaging enables spatially resolved, label-free detection of metabolites in tissue and culture systems, providing insight into their metabolic landscape and spatial distribution. However, conventional approaches often lack the spatial resolution and specificity needed to investigate metabolic heterogeneity at the single-cell level, particularly in physiologically relevant models. Here, we present a single-cell ambient mass spectrometry imaging platform, enabling direct chemical mapping of metabolites at 10 μm resolution. This method integrates cell labelling, high resolution microscopy and AP-MALDI Orbitrap mass spectrometry imaging to achieve cell-type-specific metabolite profiling. To demonstrate its application, we applied this approach to glioblastoma (GBM), an aggressive adult brain tumour characterised by cellular heterogeneity, metabolic adaptation, and infiltrative growth within the tumour microenvironment. A co-culture model combining patient-derived glioblastoma invasive-margin cells with human cortical astrocytes was used to recapitulate the invasive niche. Distinct metabolic signatures emerged upon glioblastoma–astrocyte interaction, involving pathways related to nucleotide metabolism, phospholipid and sphingolipid turnover, and tryptophan and tyrosine metabolism. These findings suggest cell-type-specific metabolic activity and potential intercellular metabolic interplay. Overall, this workflow offers a broadly accessible and robust approach for investigating metabolic heterogeneity at cellular resolution, enabling insights into metabolic interactions of heterogenous cell types in both disease and non-disease settings.

## Introduction

Bulk metabolomics has enabled insights into global metabolic trends by analysing populations of thousands to millions of cells simultaneously.^1^ Such approaches often mask subtle but biologically important heterogeneity within and between individual cells.^2^ To overcome this limitation, a variety of single-cell metabolomics techniques have been developed to investigate cellular metabolism at the single-cell level, including secondary ion mass spectrometry (SIMS), matrix-assisted laser desorption/ionisation mass spectrometry (MALDI-MS), electrospray ionisation MS (ESI-MS), and dielectric barrier discharge ionisation (DBDI).^2,3^ Among these, imaging mass spectrometry methods such as SIMS and MALDI are particularly valuable, as they provide label-free metabolite mapping.^4^ SIMS offers exceptional spatial resolution and sensitivity for small molecules, but extensive fragmentation can limit detection of larger metabolites and lipids.^5^ MALDI provides a softer ionisation that enables the detection of a broad range of biomolecules, including metabolites, lipids, and peptides, making it particularly well-suited for single-cell metabolic analysis. Unlike vacuum MALDI, AP-MALDI-MS operates under ambient conditions, reducing sample preparation constraints and better preserving volatile metabolites such as short-chain fatty acids, volatile organic acids and small polar compounds that may be lost under vacuum conditions. Although AP-MALDI-MS typically produces lower ion intensities, its combination with a high-resolution Orbitrap mass analyser offering high mass accuracy and resolving power mitigates this limitation.^6–8^ This integration enables annotation of isobaric metabolites through the high mass resolving power of AP-MALDI–Orbitrap workflows, making it particularly well-suited for spatially resolved metabolomic analyses.^3,9,10^

Recent innovations in spatial metabolite imaging have advanced the ability to visualise metabolic activity with cellular precision. Workflows such as SpaceM, HT SpaceM and the approach developed by Zhang et al. have demonstrated the feasibility of imaging metabolites at single-cell resolution.^11–13^ Despite these advances, a critical limitation remains: most existing approaches lack the ability to precisely identify individual cell types within heterogeneous samples. This constraint limits their application to co-culture and tissue systems where multiple, metabolically distinct cell populations interact. Bridging this gap by integrating spatially resolved metabolite mapping with robust cell-type identification would enable direct investigation of intercellular metabolic communication within complex microenvironments, such as tumour niches composed of cancer cells, and diverse supportive stromal cells.

To address the unmet need, we developed a microscopy-guided single-cell AP-MALDI-mass spectrometry imaging (AP-MALDI-MSI) workflow, integrated with a high-resolution Orbitrap mass analyser, to enable spatially resolved metabolomic profiling at the single-cell level. Given the highly invasive nature of glioblastoma and its interactions with surrounding astrocytes that contribute to recurrence, delineating metabolic crosstalk at the single-cell level is essential to uncover potential vulnerabilities. To retain information regarding cell identity within mixed populations, we incorporated fluorescence cell-labelling for human cortical astrocytes and patient-derived cancer cells. This approach supported the analysis of complex co-culture systems while maintaining single-cell specificity and spatial context. To our knowledge, this represents the first demonstration of glioblastoma–astrocyte metabolic crosstalk captured at single-cell resolution using AP-MALDI-MSI, a versatile framework for interrogating heterogeneous intercellular metabolic interactions in disease and non-disease states.

## Experimental Section

### Cell Culture

Patient-derived GBM cells used for single-cell mass spectrometry analysis were isolated from the 5-aminolevulinic acid (5-ALA) fluorescence-positive invasive margin of human GBM biopsy tissue during fluorescence-guided neurosurgery.^14^ These GBM invasive margin cells, referred to as GIN 8, GIN 28, and GIN 31, correspond to individual patients. All GIN cell lines were cultured in Dulbecco’s Modified Eagle Medium (DMEM) (Sigma-Aldrich, St. Louis, MO, USA) supplemented with 10% foetal bovine serum (FBS), 1 g/L glucose, and 2 mM L-glutamine (Sigma-Aldrich). Human cortical astrocytes (HA) (ScienCell Research Laboratories, Cat. No. 1800) were cultured in Astrocyte Medium (ScienCell, Cat. No. 1801) following the manufacturer’s instructions. Both GBM and astrocyte cultures were maintained at 37 °C, in a humidified atmosphere with 5% CO₂ and 90% relative humidity. For co-culture experiments, both cell types were maintained in Astrocyte Medium (ScienCell, Cat. No. 1801) to support optimal astrocyte and cancer cell viability.

GIN cells were genetically labelled with eGFP using a lentiviral construct expressing eGFP under the control of the EF1-α promoter. Viral particles were introduced at a multiplicity of infection (MOI) of 50–100, depending on the cell line, with polybrene used to enhance transduction efficiency. Human cortical astrocytes were labelled with CellTrace™ Violet Cell Proliferation Kit (Invitrogen™, Thermo Fisher Scientific, Cat. No. C34557) at a dilution of 2 μL per 5 mL of PBS, following the manufacturer’s protocol.

### Sample Preparation

Cells were cultured in 8-well μ-Slides with removable chambers (Ibidi, cat. no. 80826), pre-coated with poly-L-lysine (PLL; ScienCell, cat. no. 0413) at a concentration of 15 μg/mL to enhance cell adhesion. For monoculture conditions, 10,000 cells were seeded per well. In co-culture conditions designed to mimic the GBM invasive margin, 3,000 GIN cells and 7,000 HA cells were seeded onto each well. On day 5 of culture, cells were washed with phosphate-buffered saline (PBS), fixed with 4% PFA for 15 minutes, and subsequently rinsed with PBS followed by sterile water. The slide was then desiccated for 15 minutes to remove residual moisture prior to analysis. The chosen seeding ratio (30% GIN to 70% HA) reflects proportions reported in human GBM tissue at the invasive margin.^15,16^

### Pre- and Post-Imaging Microscopy

Microscopy was performed on the same day as AP-MALDI-MSI analysis using a Nikon Eclipse Ti inverted microscope equipped with a Plan Fluor 10×/0.30 NA objective. Brightfield and fluorescence images were acquired using standard acquisition software. eGFP fluorescence (Ex ∼488 nm, Em ∼509 nm) and CellTrace™ Violet (CTV; Ex ∼405 nm, Em ∼450 nm) were used to visualise GBM and astrocyte populations, respectively. Post-ablation images were acquired after AP-MALDI-MSI to visualise laser ablation marks, which were then used for image alignment and overlay to aid in single-cell ROI identification. Imaging parameters were kept consistent across all samples, and analyses were complemented by using ImageJ software.

### Matrix Application

Matrix application was performed using a SunCollect automated pneumatic sprayer (SunChrom, Friedrichsdorf, Germany) to uniformly coat the surface of fixed GBM and astrocyte cell samples. For matrix optimisation, three matrices were investigated: 2,5-dihydroxybenzoic acid (DHB) prepared at 7.5 mg/mL in 70:30 (v/v) acetonitrile:water (ACN:H₂O), α-cyano-4-hydroxycinnamic acid (CHCA) at 5 mg/mL in 70:30 (v/v) ACN:H₂O, and 5mg/mL 9-aminoacridine (9AA) dissolved in 70:30 (v/v) methanol (MeOH):water. Matrices were applied in 18 layers, beginning with a flow rate of 20 μL/min for the first layer, followed by 40 μL/min for the second layer, and 60 μL/min for all subsequent layers. The nozzle-to-sample distance was maintained at 2 mm, with a spraying velocity of 800 mm/min and a drying time of 2 seconds between each layer to ensure uniform crystallization and matrix coverage.

### AP-MALDI-MSI and Data Processing

Samples were analysed using an AP-MALDI UHR (MassTech Inc., Columbia, MD, USA) ion source coupled to a Q Exactive Orbitrap mass spectrometer (Thermo Fisher Scientific, Hemel Hempstead, UK). Instrument parameters for the AP-MALDI source were configured via the Target-ng software (MassTech Inc.). The source was operated in raster mode with a laser spot size of 10 μm, laser energy set to 4.5%, and a laser repetition rate of 2.5 kHz. The ion transfer capillary temperature was maintained at 400 °C and the spray voltage was set to 2.5 kV.

Mass spectrometric acquisition was performed in both positive and negative ion modes over an *m/z* range of 75–1050. The mass resolving power was set to 140,000 at m/z 200. The automatic gain control (AGC) target was 4 × 10⁶, with a maximum injection time of 400 ms, and a scan rate of 1 scan/s.

Mass spectrometry imaging data was acquired using an AP-MALDI system (MassTech Inc., Columbia, MD, USA), and mass spectral data were collected using Xcalibur software (Thermo Fisher Scientific). Raw data files were converted to the imzML and ibd formats using ProteoWizard and the MT imzML Converter (MassTech Inc.), enabling compatibility with downstream analysis platforms.^17^

Preprocessing and normalisation were performed using the *Cardinal* package in R.^18^ Total ion current (TIC) normalisation was applied to correct for overall intensity variation. Baseline correction was carried out by interpolating a baseline from local minima in the spectra.^19^ Peak picking was performed by estimating a signal-to-noise ratio (SNR) of 5, calculated from the mean absolute deviation of wavelet-convolved spectra. Peaks were then aligned across spectra using a 5 ppm *m/z* tolerance.

MS images were first generated as ion intensity maps for selected *m/z* values in *Cardinal* package in R. These were then aligned with pre-MS fluorescence microscopy images and post-ablation brightfield images to identify regions of interest (ROI) corresponding to single cells. Ion intensity values were subsequently extracted from these ROIs for downstream statistical analysis.

### AP MALDI Data Analysis

To visualise global metabolic variation across single-cell monoculture samples, a heatmap was generated using the pheatmap package in R (version 4.3.1). The dataset included all identified human metabolites, yielding a total of 88 annotated lipids and metabolites. Prior to visualisation, intensity values were log_10_-transformed and subjected to z-score normalisation across each feature to centre the data and facilitate comparative interpretation. Samples were arranged in a predefined group order without clustering to preserve the experimental structure, while metabolites were clustered using unsupervised hierarchical clustering based on Euclidean distance to reveal patterns of co-abundance.

Principal Component Analysis (PCA) was conducted to visualise variance in metabolite abundance across experimental groups and to identify underlying patterns in the data. Prior to PCA, metabolite intensity values were log₁₀-transformed to reduce right skewness, stabilise variance, and improve the normality of the data distribution. Pareto scaling was subsequently applied to moderate the dominance of high-intensity ions and preserve relative variance among metabolites.^20,21^ This approach has recently been shown in Orbitrap-based imaging datasets to enhance the interpretability of multivariate analyses compared with unscaled data.

To classify samples and identify discriminative features, a Random Forest (RF) model was implemented using the *caret* and *randomForest* packages in R (version 4.3.1). Model training involved 10-fold cross-validation, repeated three times to ensure robust performance evaluation. Each model comprised 500 trees, with the number of variables randomly sampled at each split optimised automatically by cross-validation. Reproducibility was maintained by setting a fixed random seed (set.seed = 123). The classification model was evaluated based on the area under the receiver operating characteristic curve (AUC-ROC), providing a measure of model discrimination. Feature importance was assessed using both Mean Decrease in Accuracy (MDA) and Mean Decrease in Gini index (MDG) derived from a Random Forest classifier.^22,23^ Features with MDA >1 and MDG >0 were considered significant contributors to class separation consistent with previous applications of Random Forest in metabolomics. These thresholds were empirically supported by the distribution of feature scores within our dataset and validated using permutation testing (100 iterations, *rfPermute* package).

Univariate statistical analysis was performed using MetaboAnalyst 6.0.^24^ To compare metabolite levels between monocultured GIN and HA cells, an unpaired, non-parametric Mann–Whitney U test was applied. This test was chosen due to the non-normal distribution of the data and differences in group variance, as assessed by Levene’s test. Benjamini–Hochberg correction was used to control the false discovery rate (FDR) associated with multiple testing. For co-culture experiments, two comparisons were performed: (1) HA in co-culture vs. HA in monoculture, and (2) GIN in co-culture vs. GIN in monoculture, using data across 10 replicates (i.e. 10 individual cells per cell line). Additionally, pairwise comparisons were conducted for each individual cell line pairing using the Kruskal–Wallis test. For all comparisons, log₂ fold changes (log₂FC) were calculated to determine the direction and magnitude of change. Metabolites were considered significantly altered if |log₂FC| > 1.5 and FDR-adjusted p < 0.05.

### Metabolite Identification

Metabolite and lipid identification was conducted using a molecular formula prediction tool developed by the School of Pharmacy, University of Nottingham.^25^ Mass spectral features from both positive and negative ion modes were screened against the Human Metabolome Database (HMDB) and LIPID MAPS using a mass tolerance of ±5 ppm.

### Liquid Chromatography–Mass Spectrometry (LC-MS) analysis

LC-MS/MS was performed to validate and compare metabolites identified by AP-MALDI-MS, providing confirmation and increasing confidence in metabolite assignments. GIN 8, and GIN 31 cells were cultured in T75 flasks to a density of approximately 5 × 10⁵ cells on the day of extraction. Cells were fixed with 4% PFA for 15 minutes at room temperature, followed by sequential washes with PBS and sterile water to remove residual fixative and salts. To quench metabolism and initiate extraction, cells were treated with 500 μL of ice-cold (−20 °C) LC-MS-grade MeOH. Metabolites were extracted by manually scraping the cells and collecting the supernatant. The cell extracts were then incubated on a thermomixer at 4 °C and 2,000 rpm for 1 hour to facilitate metabolite release. Following incubation, samples were centrifuged at 13,000 × g for 5 minutes at 4 °C to separate the cellular debris from the metabolite-containing supernatant. The MeOH supernatants were collected and evaporated using a vacuum concentrator (SpeedVac; Thermo Fisher Scientific) at room temperature.

Dried extracts were then reconstituted in 70 μL of LC-MS-grade MeOH, followed by a second centrifugation at 13,000 × g for 10 minutes at 4 °C to remove any remaining particulate matter. The resulting supernatants were carefully transferred to LC-MS vials and stored at −80 °C until analysis.

Prior to LC-MS analysis, quality control (QC) samples were prepared by pooling 5 μL aliquots from each individual extract to assess system stability and reproducibility across the run.

Samples were analysed on a Q Exactive Orbitrap mass spectrometer (Thermo Fisher Scientific, Hemel Hempstead, UK) equipped with a HESI-II electrospray ionisation source operating in positive and negative ion switching mode. Full-scan MS data were acquired over an *m/z* range of 200–2000 at a resolution of 140,000 at m/z 200. Data-dependent MS/MS (ddMS²) acquisition was conducted on pooled QC samples using a top-5 method at a resolution of 17,500 at m/z 200 with stepped normalised collision energies of 20, 30, and 40.

Chromatographic separation was achieved using a ZIC-pHILIC column (150 × 4.6 mm², 5 μm particle size) maintained at 45 °C on a Dionex UltiMate 3000 LC system (Thermo Fisher Scientific, Hemel Hempstead, UK). Elution was performed using a solvent system consisting of phase A (10mM ammonium carbonate in water) and phase B (acetonitrile). A linear gradient from 80% to 5% of mobile phase B was applied over 15 minutes, followed by a return to 80% B over 2 minutes and a 7-minute re-equilibration at 80% B, giving a total run time of 24 minutes. The flow rate was 300 μL/min, the injection volume was 10 μL, and samples were maintained at 4 °C throughout.

Raw LC-MS data were processed using *Compound Discoverer* 3.3 SP2 (Thermo Fisher Scientific, Hemel Hempstead, UK). An untargeted analysis workflow was employed for peak detection, alignment, and preliminary compound annotation. Accurate mass values were matched against the HMDB and mzCloud spectral library (https://www.mzcloud.org/). For features with MS/MS spectra, compound identification was supported by spectral matching in mzCloud. To further support identification confidence, analytical standards were run alongside biological samples. These standards were used to confirm the retention times and MS/MS fragmentation patterns of selected metabolites, enabling assignment of Schymanski confidence levels.^26^

LC–MS/MS of monocultured cells identified 33 metabolites also detected by AP-MALDI-MSI. This overlap, while obtained under different culture conditions, supports the structural assignments of these metabolites in AP-MALDI-MSI datasets.

## Results and Discussion

### AP-MALDI-MSI Single-Cell Metabolomics Workflow

To enable single-cell level metabolic profiling, we developed and implemented a streamlined workflow integrating microscopy-guided cell identification with AP-MALDI-MSI. As illustrated in **Figure 1A–D**, cells were seeded onto chamber slides and fixed using 4% paraformaldehyde (PFA) to preserve spatial architecture and metabolite integrity. Following fixation, brightfield microscopy was used to capture cell morphology and guide image co-registration. After matrix application, AP-MALDI-MSI was performed using high-resolution mass spectrometry at 10 µm spatial resolution.

**Figure 1.**
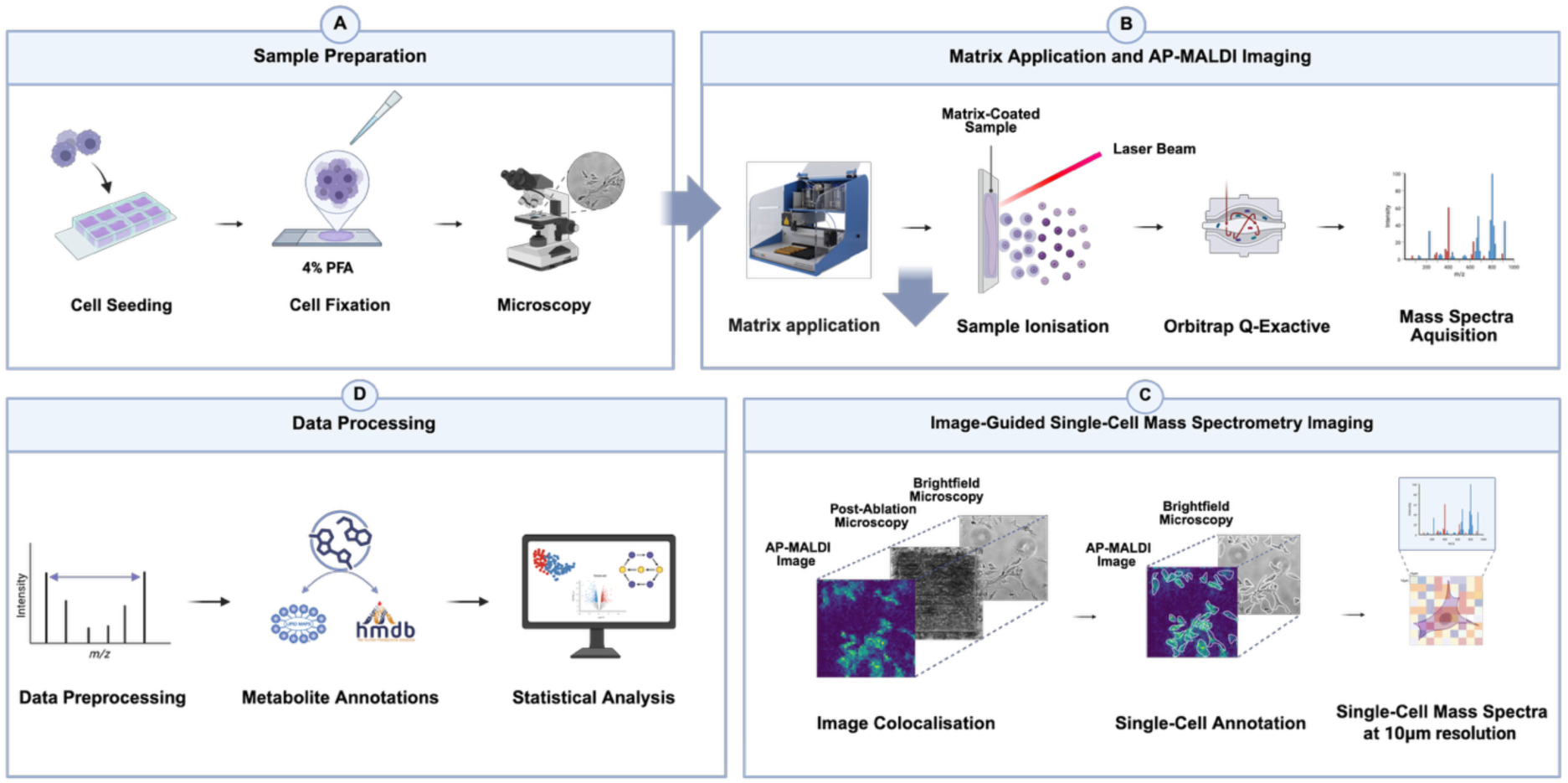
Workflow for AP-MALDI-MSI based single-cell metabolomics. **(A)** Cells are seeded onto chamber slides, fixed using 4% PFA, and imaged using brightfield microscopy for cell identification. **(B)** Matrix is applied to the sample, followed by laser ablation and ionisation to acquire AP-MALDI-MSI images. **(C)** Image registration is performed to align brightfield microscopy, ablated region, and AP-MALDI-MSI data, enabling extraction of single-cell metabolomes. **(D)** Data processing steps for generating and analysing single-cell metabolic profiles.

Our workflow incorporated microscopy, both pre- and post-ablation, to precisely align ablated regions with microscopy images for single-cell identification. To integrate AP-MALDI-MSI data at the single-cell level, MSI datasets were processed and spatially registered in R using the *Cardinal* package, with adaptations for coordinate alignment and pixel-level intensity extraction from single cells. Preprocessing included total ion current normalisation, local-minima baseline reduction, and peak alignment at 5 ppm tolerance. Coordinate mapping between microscopy and MSI datasets was optimised to account for AP-MALDI-MSI discrete ablation geometry, enabling precise ion extraction from individual cells. Extracted features were subsequently annotated through molecular formula prediction using in-house software developed at the School of Pharmacy, University of Nottingham, based on accurate mass and isotopic pattern matching.^25^ Although this approach has previously been applied in OrbiSIMS datasets, this study represents the first demonstration of its integration within AP-MALDI-MSI data for single-cell metabolomic analysis.

### Comparative Analysis of AP-MALDI-MSI Matrices for Single-Cell Metabolite Profiling

We first evaluated matrix selection for single-cell metabolomics using AP-MALDI-MSI by investigating three matrices: 2,5-dihydroxybenzoic acid (DHB), α-cyano-4-hydroxycinnamic acid (CHCA) and 9-aminoacridine (9AA). As shown in **Figure 2A**, DHB enabled clear identification of the matrix peak at *m/z* 155.0338 and a cell-specific metabolite peak at *m/z* 230.1587, corresponding to FA 28:7;O3. The spatial distribution of this ion delineated single-cell morphology, confirmed by overlaying the brightfield image with reduced opacity onto the AP-MALDI-MSI ion image. Representative mass spectra from HA cells in each mode are shown, highlighting DHB matrix-associated peaks—C₇H₇O₄⁺ in positive mode and C₇H₅O₄⁻ in negative mode. We performed parallel analyses using CHCA and 9AA to compare metabolite coverage and chemical class representation as shown in **Figure 2B–C**. While each matrix generated distinct profiles, CHCA and DHB exhibited the greatest overlap in detected features, followed by DHB and 9AA. These results are consistent with previous studies reporting matrix-dependent selectivity in metabolite detection.^27^ The chemical class distribution revealed further matrix-specific trends; DHB and CHCA primarily enriched amino acids and their derivatives, whereas 9AA preferentially ionised sterol lipids, **Figure 2D**. Importantly, DHB yielded the highest number of metabolite features overall (*n* = 412), compared to CHCA (*n* = 236) and 9AA (*n* = 230). To enable broad metabolomic coverage encompassing both small polar metabolites (e.g., amino acids) and diverse lipid classes at the single-cell level, DHB was selected as the matrix for subsequent comparisons between glioblastoma invasive margin cells (GIN) and astrocytes (HA). This systematic optimisation provides the first direct comparison of matrix-dependent metabolite coverage at the single-cell scale in ambient conditions, offering practical guidance for future single-cell workflows. Having established DHB as the optimal matrix, we next compared single-cell AP-MALDI-MSI with conventional bulk LC-MS to characterise differences in metabolite coverage and to evaluate whether cross-platform overlap can be used to increase confidence in metabolite assignments.

**Figure 2.**
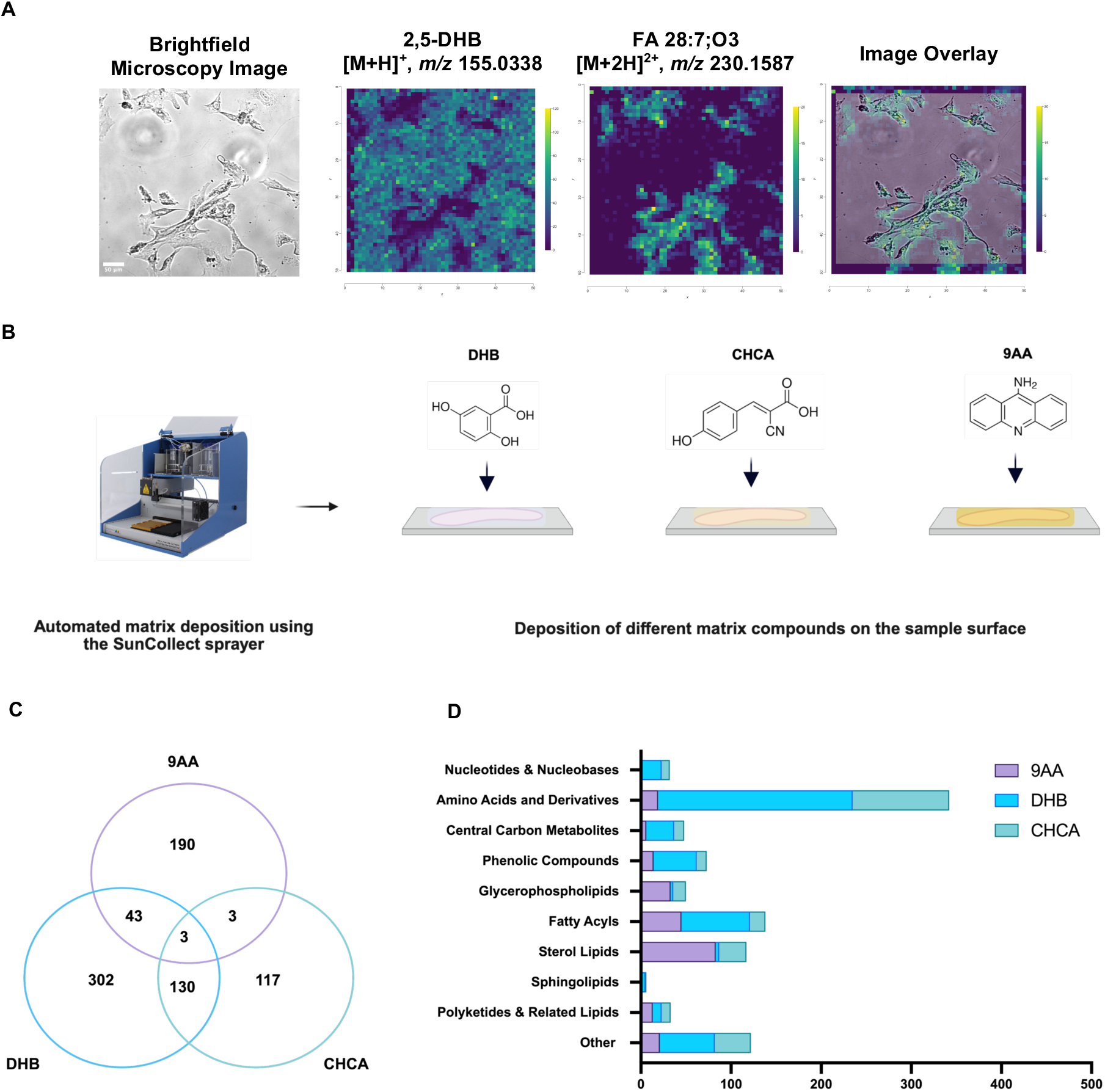
Comparative analysis of metabolite detection in human cortical astrocytes using AP-MALDI-MSI with different matrices. **(A)** Brightfield microscopy, magnification x10 and AP-MALDI-MSI ion images representing DHB matrix peak at *m/z* 155.0338, followed by cell specific marker FA 28:7;O3 at *m/z* 230.1587, and AP-MALDI-MSI ion image and brightfield microscopy image overlay to illustrate spatial metabolite distribution in HA along with representative mass spectra acquired in both positive and negative ion modes. **(B)** Schematic of automated matrix deposition using the SunCollect sprayer and chemical structures of DHB, CHCA, and 9AA. **(C)** Venn diagram showing the overlap of metabolites detected using AP-MALDI-MSI with DHB, CHCA, and 9AA matrices. **(D)** Bar graph comparing the number and class of metabolites detected with different matrices across various compound categories.

When comparing HA cells analysed by AP-MALDI-MSI with bulk populations of approximately 5×10^5^ cells profiled by LC-MS as shown in **Figure 3A**, we observed higher number of metabolite annotation in bulk LC-MS method compared to single-cell AP-MALDI-MSI method, **Figure 3B-C**. Nevertheless, the single-cell AP-MALDI-MSI workflow yielded several hundred annotated metabolites, representing a remarkably high number relative to sample size. We observed limited overlap between platforms, with 33 shared metabolites detected by both, **Figure 3B**. LC-MS predominantly captured polar metabolites, including amino acids and derivatives and central carbon metabolites, **Figure S1**, whereas AP-MALDI-MSI detected amino acid and lipid species, as shown earlier in **Figure 2D**. To improve identification confidence, we acquired LC-MS/MS annotating structural assignments for metabolites initially observed by AP-MALDI-MSI, **Table S1**.^28^ The limited overlap of metabolites between two techniques observed is consistent with previous reports, reflecting intrinsic differences in ionisation efficiency, detection sensitivity and chromatographic separation associated with the LC method.^29^

**Figure 3.**
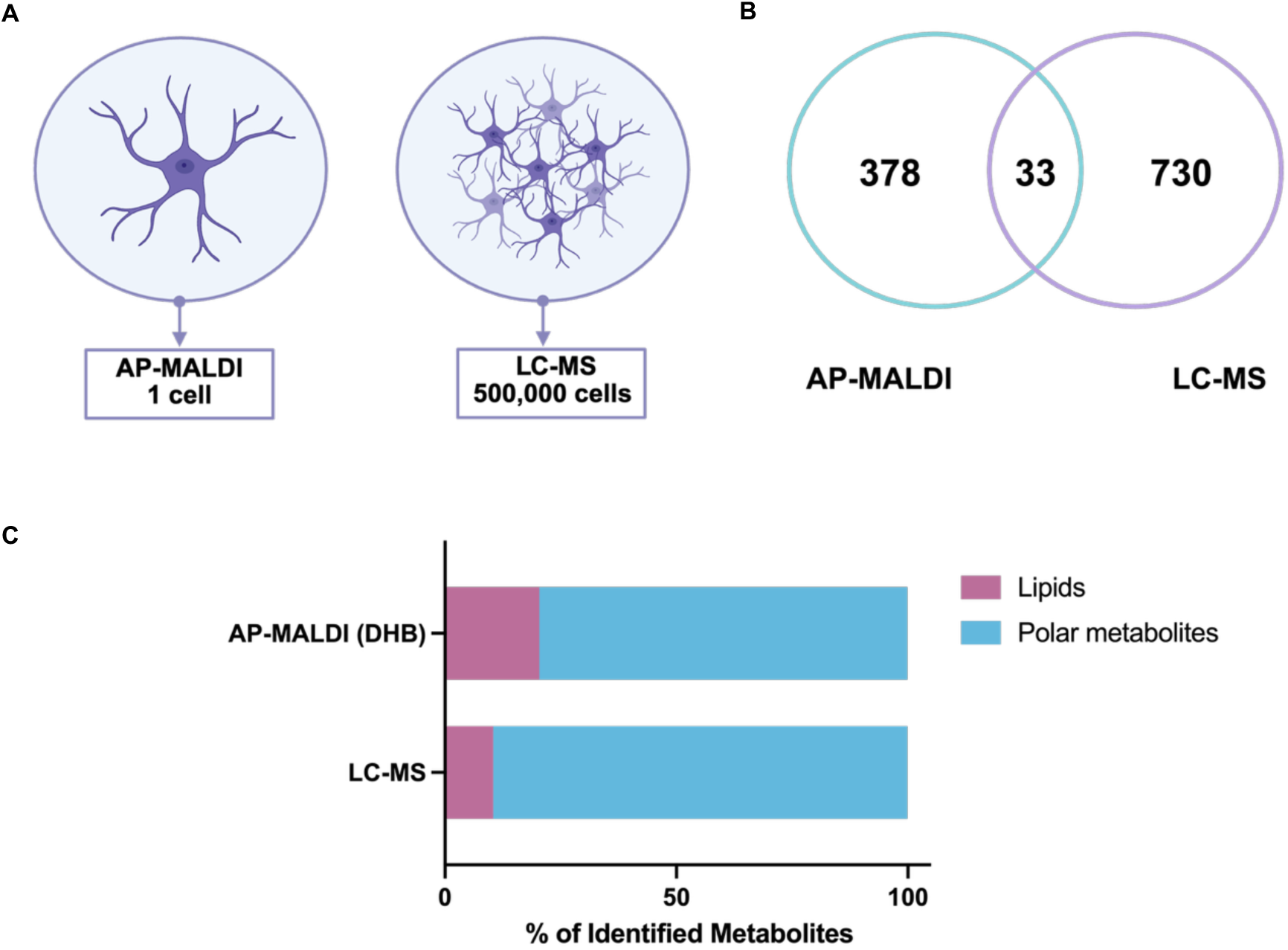
Comparative overview of AP-MALDI-MSI and LC-MS metabolite detection. **(A)** Schematic representation of sample input: AP-MALDI-MSI analysis of a single astrocyte versus LC-MS analysis of 5×10^5^ astrocytes. **(B)** Venn diagram showing the number of metabolites uniquely detected by LC-MS (*n* = 730) or AP-MALDI-MSI with DHB matrix (*n* = 378), and the overlap between platforms (*n* = 33). **(C)** Distribution of single cell identified metabolites by chemical class, highlighting differences in lipid (violet) and polar metabolite (blue) coverage.

Overall, these results establish a robust single-cell AP-MALDI-MSI workflow capable of broad, untargeted metabolite profile from single cells under ambient conditions, spanning small polar metabolites to multiple lipid subclasses. While bulk LC-MS remains advantageous for broader polar metabolite coverage, our cross-platform evaluation demonstrates that single-cell AP-MALDI-MSI uniquely enables cell-specific metabolic investigation with spatial context, which is an analytical capability not achievable by bulk methods. This foundational optimisation provides the analytical basis for subsequent studies to investigate metabolic interactions between different cell types in co-culture.

### Metabolite Profiling Reveals Cell-Specific Signatures for Astrocyte and GBM Single-Cell Annotations

We then performed a comparative analysis to investigate metabolic differences between human astrocytes (HA) and glioblastoma invasive margin cells (GIN) using AP-MALDI-MSI (**Figure S2A–B**). GIN cells exhibited broader intra-group variance than HA cells, as indicated by their wider distribution along PC1 (24.66%) and PC2 (23.88%), reflecting greater metabolic heterogeneity within the invasive tumour population. This is consistent with the pronounced heterogeneity reported in invasive glioblastoma populations, where metabolic plasticity supports adaptation and therapeutic resistance.^30,31^ In contrast, the tight clustering of HA cells reflects the more uniform metabolic profile of HA, whose core functions in glucose metabolism, neurotransmitter cycling, and energy support are conserved across individual cells.^32^

To further investigate metabolite-level variation between GIN and HA cells, we generated a heatmap of all annotated metabolites and lipids, identified as known human compounds using the HMDB and LIPID MAPS database, **Figure 4**. Hierarchical clustering based on Euclidean distance was applied to reveal patterns of co-abundance and cell-to-cell variability within each population. This visualisation revealed pronounced metabolic heterogeneity among GIN cells, characterised by diverse intensity patterns across individual cells from the same population. In contrast, HA cells exhibited a more uniform and tightly clustered metabolite profile, consistent with lower intra-group variability. These findings highlight the advantage of single-cell metabolomics in capturing cell-to-cell metabolic diversity that would be obscured in bulk analyses—thus providing important insights into the heterogeneity of tumour-associated metabolic reprogramming. While single-cell transcriptomic and genomic approaches are comparatively more advanced and have increasingly revealed cellular heterogeneity in tumours, single-cell metabolomics remains an emerging field.^33,34^ Our study, conducted in cell culture models, demonstrates that spatially resolved metabolic profiling at the single-cell level can uncover metabolic heterogeneity, offering a novel layer of information. This approach lays important groundwork for future applications in tissue samples, where it could further illuminate tumour biology *in situ*.

**Figure 4.**
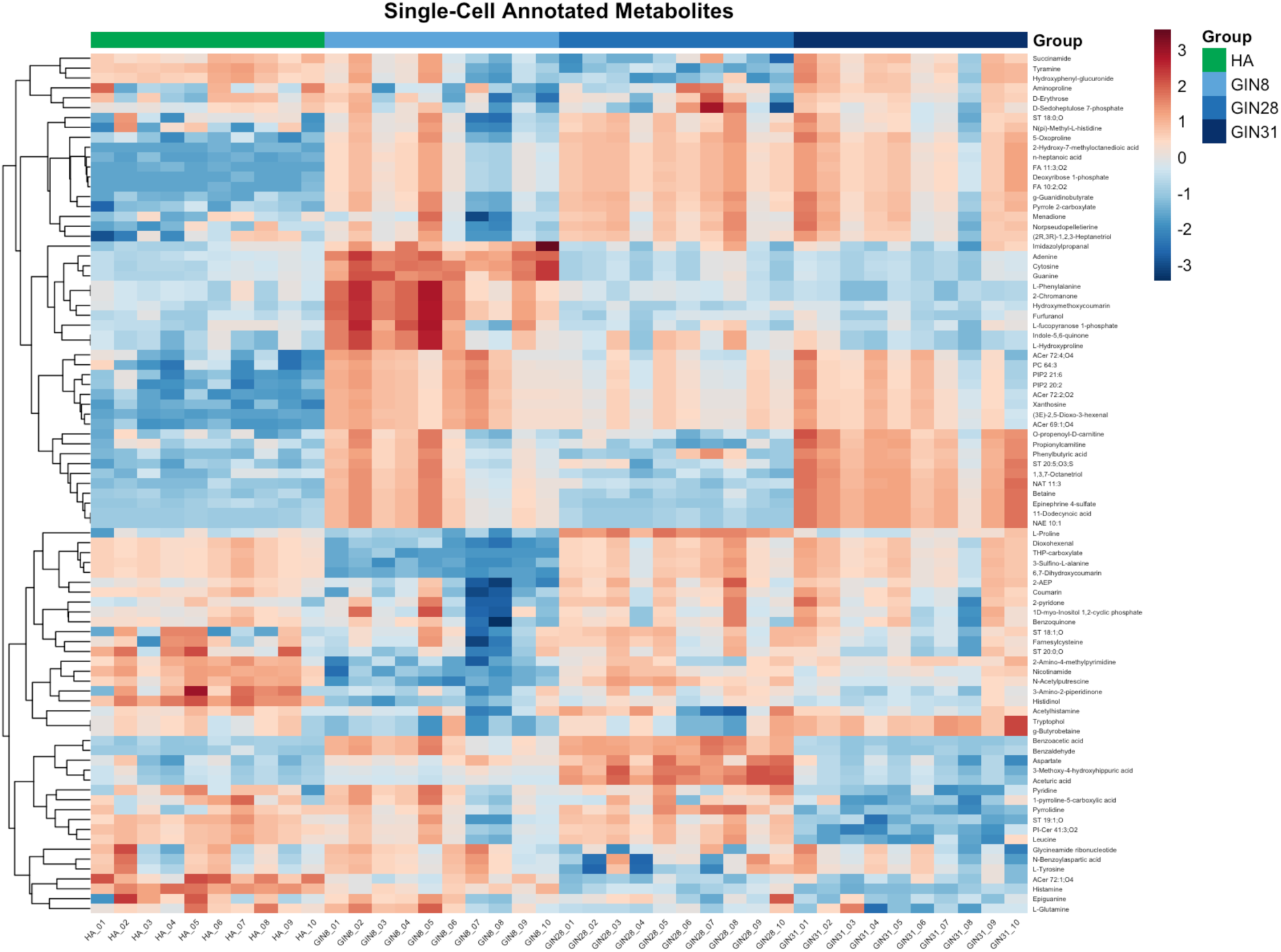
Heatmap showing the distribution of annotated metabolites in individual GIN cells (cells 8, 28, and 31, *n* = 10) and HA cells (*n* = 10). Data were z-score normalised across each metabolite to facilitate comparison between samples. Colours represent relative metabolite abundance (red = high, blue = low). Hierarchical clustering based on Euclidean distance grouped metabolites and cells by similarity

Building on these findings, we next examined how these metabolites were spatially distributed within individual cells using AP-MALDI-MSI, enabling the identification of characteristic biochemical markers.

As shown in **Figure 5A**, nucleotides and nucleobases were detected in both cell types, with guanine displaying particularly distinct spectral features. Guanine is a purine nucleobase, and elevated purine metabolism has been implicated in cell proliferation and therapeutic resistance of GBM cells, particularly through enhanced DNA repair pathways.^29^ Its consistent detection in both HA and GIN cells may reflect the fundamental role of purine turnover in brain cell function, while also hinting at metabolic pathways that GBM may exploit for survival and growth. **Figure 5B** presents the amino acids and their derivatives identified across both HA and GIN cells, among which acetylhistamine emerged as the most spatially distinct and consistently detected metabolite. Histamine and its derivatives play a role in neuromodulation, vascular regulation, and sustaining the brain microenvironment, making acetylhistamine a plausible candidate biomarker for processes shared across healthy and malignant glial cells.^29,35^ Similarly, **Figure 5C** illustrates lipid species detected in the two cell types, with FA 28:7;O3 present in both populations. Although little is known about this specific fatty acid, fatty acids are critical for membrane integrity, signalling, and energy metabolism in the brain, given the consistent detection of FA 28:7;O3 makes it a plausible marker for distinguishing and characterising brain cell populations in AP-MALDI-MSI analyses. Consequently, guanine, acetylhistamine, and FA 28:7;O3 were chosen as reference ions because they were reproducibly detected in both astrocytes and GBM cells, showed distinct spatial localisation, and are linked to key metabolic functions in brain cells. Together, these findings established a set of reproducible metabolic markers that enabled confident single-cell identification and mapping within AP-MALDI-MSI datasets. Building on this foundation, we next applied this approach to a co-culture model to explore how spatial interactions between GBM and astrocytes influence their metabolic landscape.

**Figure 5.**
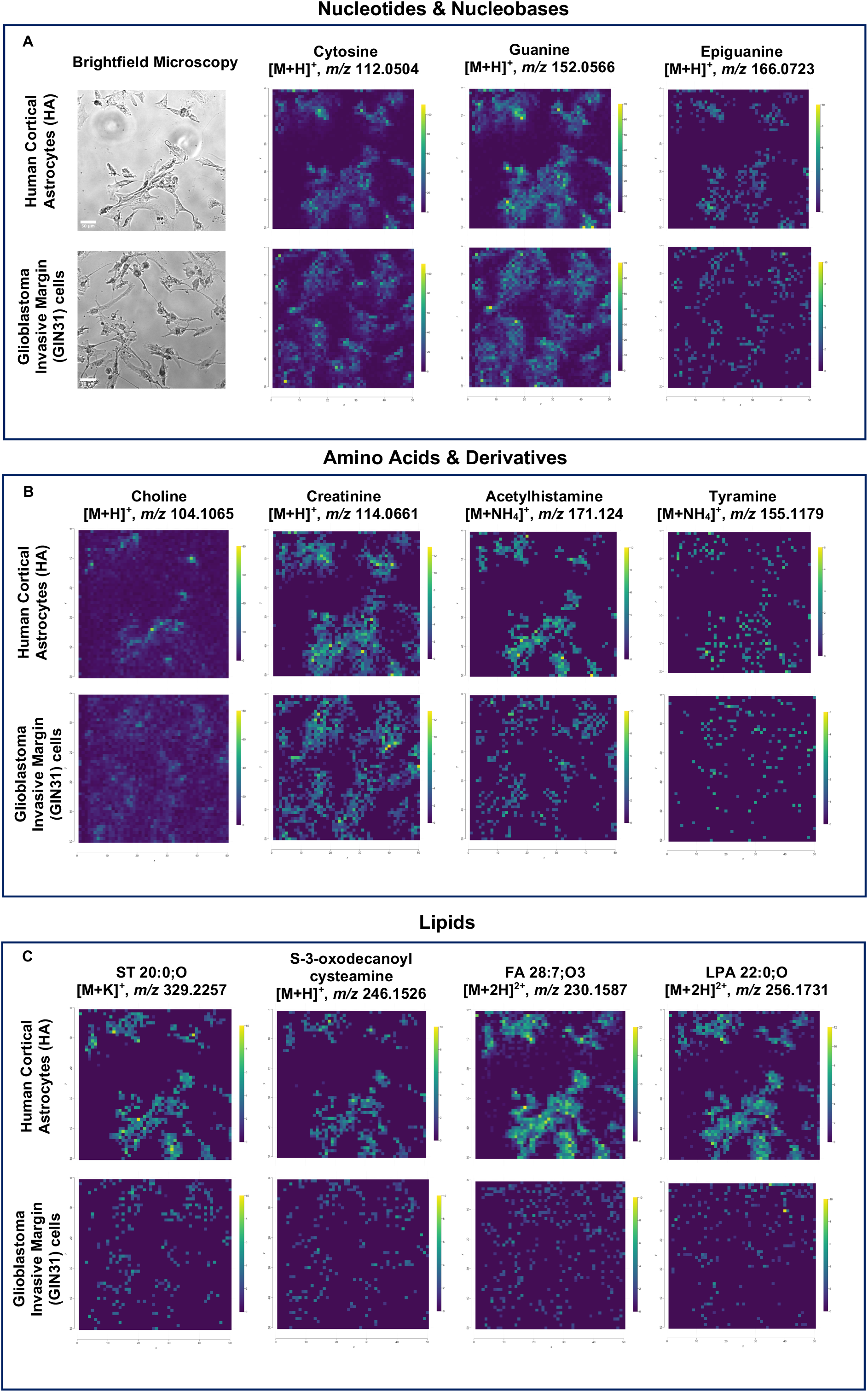
Identified metabolite and lipid classes in single cells of human cortical astrocytes and glioblastoma invasive margin cells using DHB matrix. **(A)** Panel of brightfield microscopy images of Human Astrocytes (HA) and glioblastoma invasive margin cells (GIN), alongside corresponding AP-MALDI-MSI ion images showing selected nucleotides and nucleobases. **(B)** Panel of AP-MALDI-MSI ion images displaying annotated amino acids and derivative metabolites. **(C)** Panel of AP-MALDI-MSI ion images displaying lipid annotations.

### Single-Cell Spatial Metabolomics in a Co-culture Model of Glioblastoma and Astrocytes

Having identified distinct metabolic signatures in isolated cell populations, we next applied the single-cell AP-MALDI-MSI methodology to explore metabolic crosstalk between HA and GIN cells in a co-culture setting, thus assessing feasibility for future application to primary tumour tissue composed of heterogenous disease/non-disease cell types and cellular states. Astrocytes are the most abundant glial cell type in the brain and play a key role in maintaining the tumour microenvironment.^36,37^ To mimic early-stage GBM infiltration, we established a mixed population composed of 70% astrocytes and 30% GBM patient-derived invasive margin cells.^15,16^ This model enabled direct investigation of metabolite-level interactions between the two cell types within a shared microenvironment while retaining single-cell spatial resolution, **Figure 6 A-D**.

**Figure 6.**
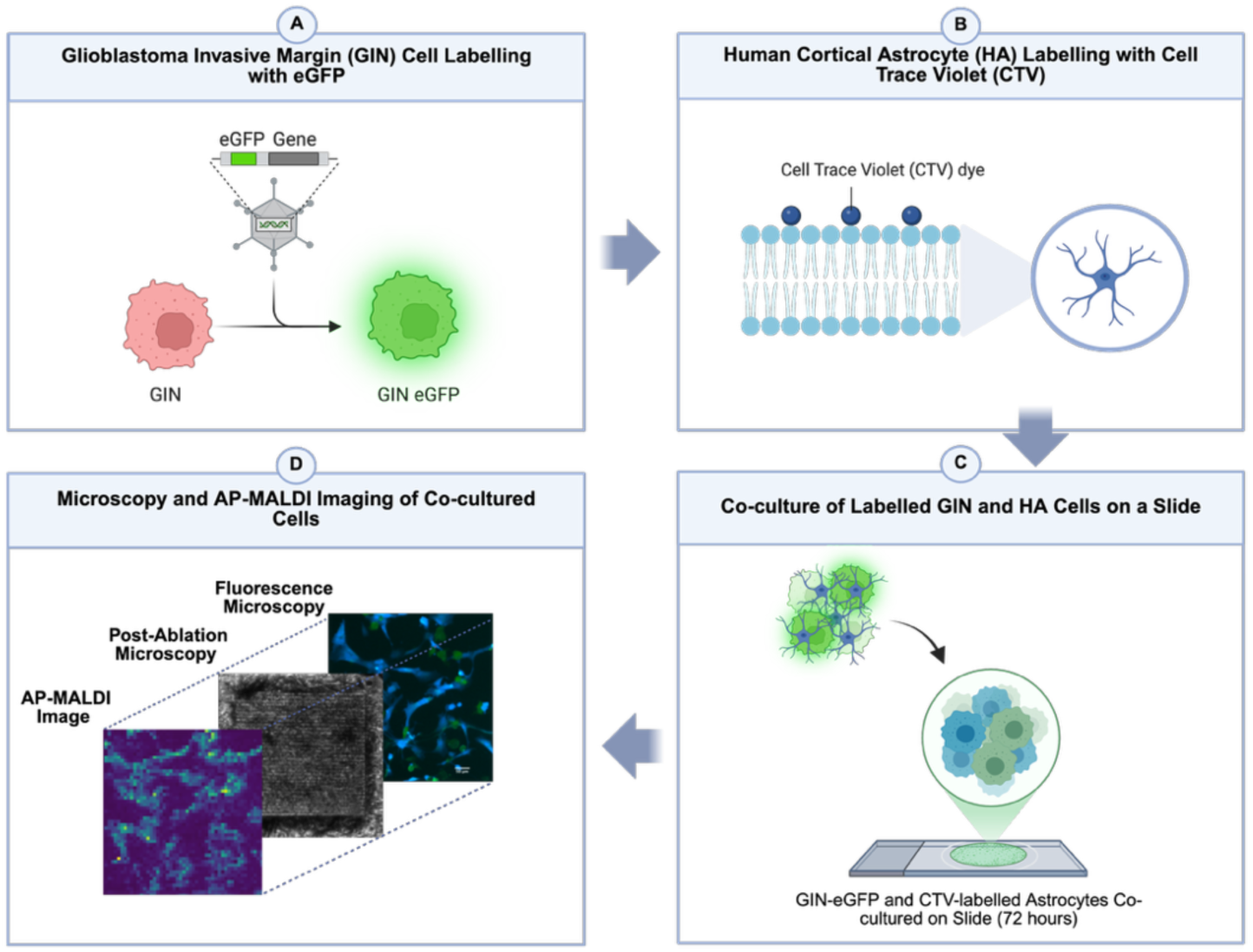
Application of single-cell AP-MALDI-MSI to investigate metabolic crosstalk in a co-culture of GIN and HA cells. **(A)** GIN cells were genetically tagged with enhanced green fluorescent protein (eGFP). **(B)** HA cells were labelled with CellTrace™ Violet dye for identification. **(C)** Co-culture of HA (CTV-labelled) and GIN (eGFP-expressing) cells was established on chamber slides. **(D)** AP-MALDI-MS image overlaid with post-ablation brightfield microscopy (10× magnification) and fluorescence microscopy images to enable single-cell analysis.

To enable precise identification of each cell type within the mixed culture, GIN cells were genetically tagged with eGFP, while HA cells were labelled with CellTrace™ Violet (CTV) membrane dye, allowing fluorescence-based single-cell discrimination. Integrating this dual-labelling strategy with AP-MALDI-MSI enabled direct correlation between fluorescence identity and metabolic profile of individual cells, establishing a robust framework for spatial single-cell metabolomics.^38^ The two cell types were co-cultured on chamber slides for five days, followed by fixation with 4% PFA. Fluorescence microscopy and post-ablation brightfield imaging were combined with AP-MALDI-MSI to extract metabolic profiles at single-cell resolution.

While previous studies have explored metabolic communication in co-culture models using mass spectrometry imaging, such as the work by Zhang et al. on Oesophageal Squamous Cell Carcinoma (ESCC) cell–fibroblast interactions using MALDI-TOF MSI, their Transwell system lacked direct cell–cell contact and offered lower mass resolution and chemical specificity. In contrast, our approach integrates direct physical co-culture with fluorescence-guided single-cell discrimination and a high-resolution mass analyser, providing a unique platform for spatially resolved metabolic profiling of different cell populations.

To gain insight into spatially distinct metabolic features associated with cell–cell interactions, we examined the distribution patterns of annotated metabolites under co-culture conditions. This exploratory analysis revealed several metabolites with markedly different localisations. As shown in **Figure 7**, metabolites such as cytosine (*m/z* 112.0504), adenine (*m/z* 136.0618), 2-aminohistamine (*m/z* 112.0868), acetylcadaverine (*m/z* 145.1335) were primarily confined within cellular boundaries, indicative of intracellular retention whereas metabolites such as 3-amino-2-piperidone (*m/z* 115.0865), and N-acetylputrescine (*m/z* 131.1179) were detected both within and along the periphery of cells. The distinction between intracellular and extracellular metabolite localisation is biologically important: intracellular metabolites reflect core metabolic processes and cell-intrinsic functional states, whereas extracellular metabolites may result from active secretion and paracrine signalling between neighbouring cells.^39–41^

**Figure 7.**
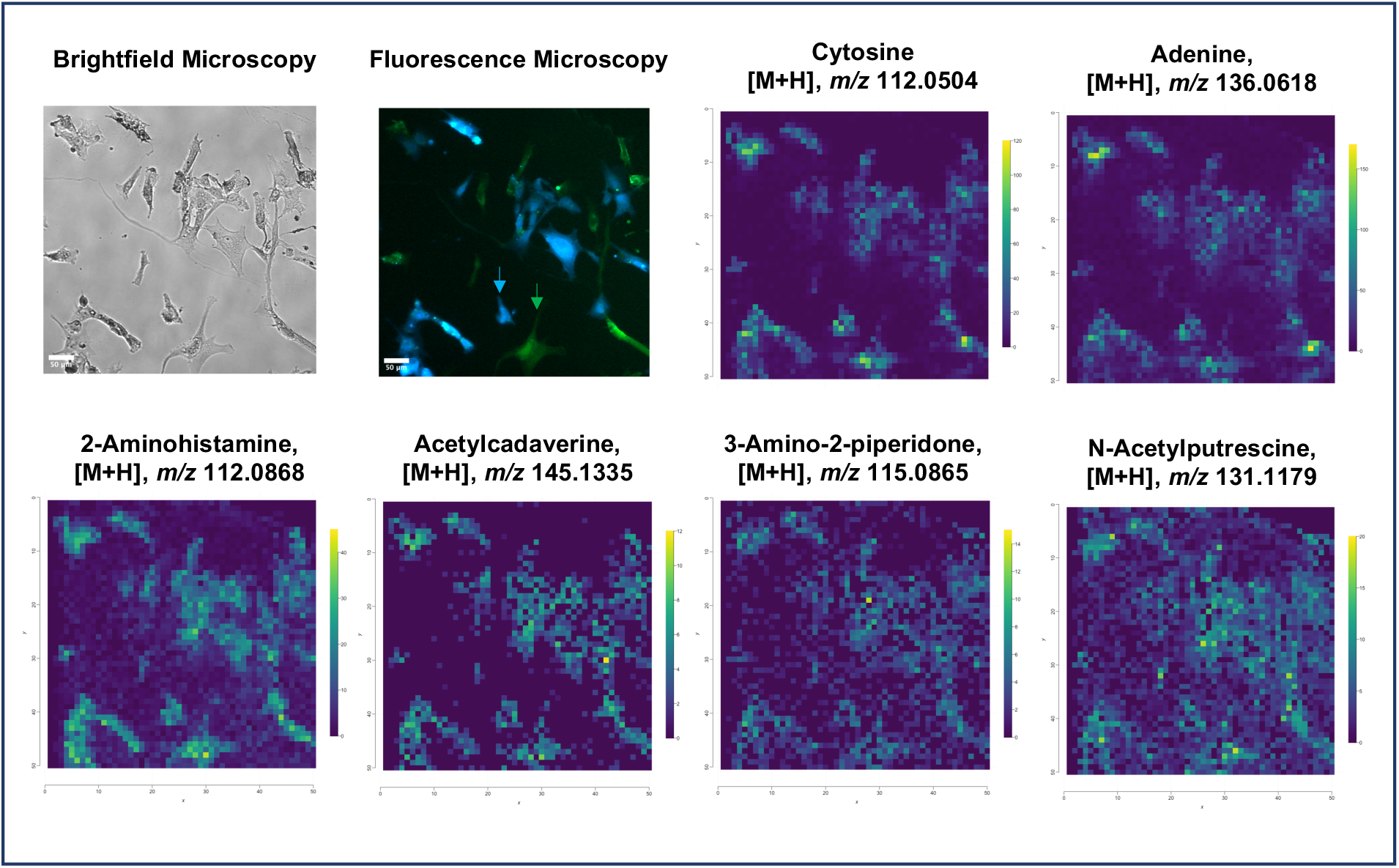
Metabolomic analysis of GIN and HA single cells from co-culture. Representative panel showing brightfield microscopy, fluorescence microscopy, and AP-MALDI images used for accurate cell annotation and extraction of individual single-cell metabolomes. In fluorescence microscopy, glioblastoma cells (eGFP⁺) are shown in green and astrocytes (CTV⁺) in blue; arrows indicate representative cells of each type.

Building on these spatial observations, we next examined how co-culture conditions influenced the overall metabolic profile of GIN cells. In **Figure 8A**, PCA illustrates the distribution of GIN cell metabolomes under co-culture and monoculture conditions. While both groups exhibit some overlap indicating shared core metabolic features, GIN cells in co-culture display a broader distribution across both principal components. This dispersion may reflect increased metabolic heterogeneity arising from intercellular interactions with astrocytes, potentially representing early metabolic adaptation or signalling-driven shifts in tumour cell metabolism.

**Figure 8.**
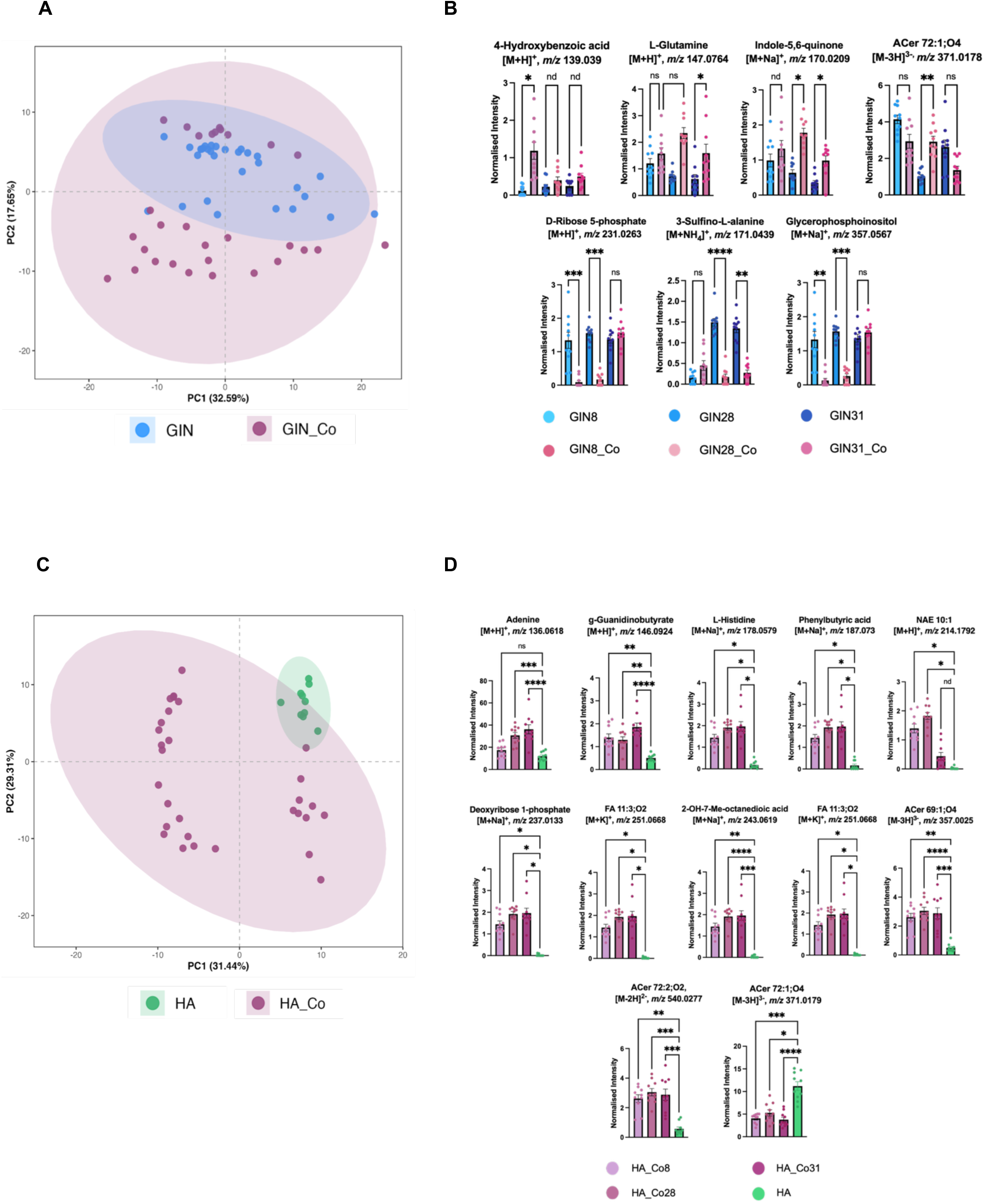
Comparison of GIN and HA cell metabolic profiles cultured in co-culture versus monoculture. **(A)** Principal Component Analysis (PCA) illustrating metabolic variation between GIN cells cultured alone (*n*=10) and those co-cultured with astrocytes (*n*=10). **(B)** significantly altered metabolites identified via univariate analysis (p < 0.05) and Random Forest feature selection. Data are presented as mean ± SEM. **(C)** PCA plot showing metabolic differences between HA cells in monoculture (*n*=10) and in co-culture with GIN cells (*n*=10). **(D)** metabolites significantly altered under co-culture conditions, identified using univariate statistics (p < 0.05) and Random Forest. Data are presented as mean ± SEM.

Subsequent statistical analysis identified metabolites and lipids that were differentially abundant in GIN cells under co-culture conditions relative to monoculture (**Figure 8B**, Table 1). D-ribose 5-phosphate (*m/z* 231.0263) and glycerophosphoinositol (*m/z* 357.0567) were significantly increased in co-cultured GIN cells, suggesting enhanced nucleotide turnover and phospholipid-related signalling in the presence of astrocytes. This metabolic shift, associated with proliferative activity, is consistent with RNA-based observations that paediatric brain cancer cells exhibit increased proliferation in response to astrocytes (McCrorie *et al*, manuscript in preparation). Similarly, L-glutamine (*m/z* 147.0764) and L-Hydroxyproline (*m/z* 170.0209) levels were elevated in co-culture, potentially reflecting metabolic rewiring of tryptophan-related pathways consistent with previous evidence that implicates tryptophan metabolism in GBM survival.^42^ Conversely, 3-sulfino-L-alanine (*m/z* 171.0439) showed a marked decrease in co-culture, which may indicate reduced sulfur amino acid metabolism or an altered redox state. As an intermediate in cysteine catabolism, 3-sulfino-L-alanine plays a key role in the biosynthesis of hypotaurine and taurine, molecules involved in cellular redox regulation.^43^ Additionally, ceramide-related lipid ACer 72:1;O4 (*m/z* 371.0178), was significantly elevated in monoculture but not in co-culture, suggesting reduced ceramide signalling or membrane remodelling in the presence of astrocytes. In the GBM microenvironment, ceramide accumulation has been associated with altered cell–cell signalling and may influence inflammatory responses and glial activation, highlighting their potential role in modulating tumour–astrocyte interactions.^44^

**Table 1.**
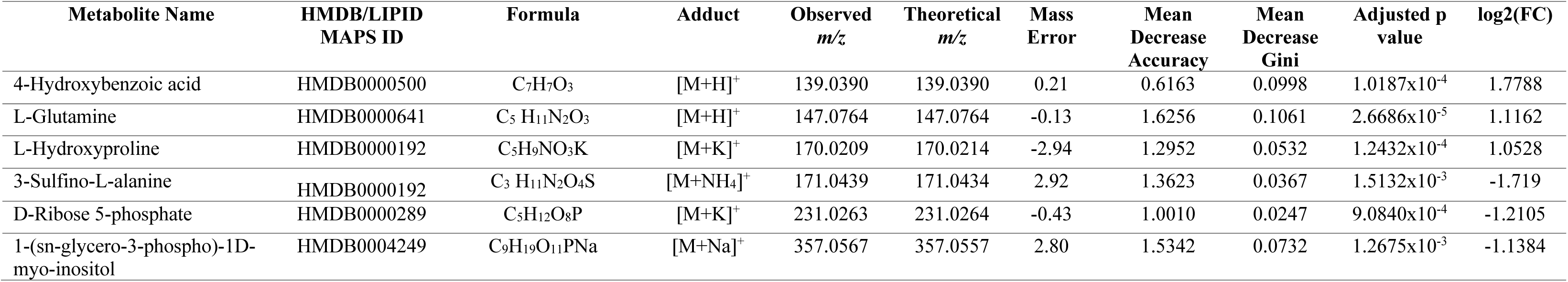
Significant metabolites identified in GIN co-culture (*n*=10) with HA (*n*=10). Metabolites were detected using AP-MALDI mass spectrometry with DHB matrix. Metabolites were selected using variable importance measures from random forest analysis (Mean Decrease Accuracy, Mean Decrease Gini), adjusted *p*-values from statistical testing, and log₂ fold change (log₂FC) values.

**Table 2.**
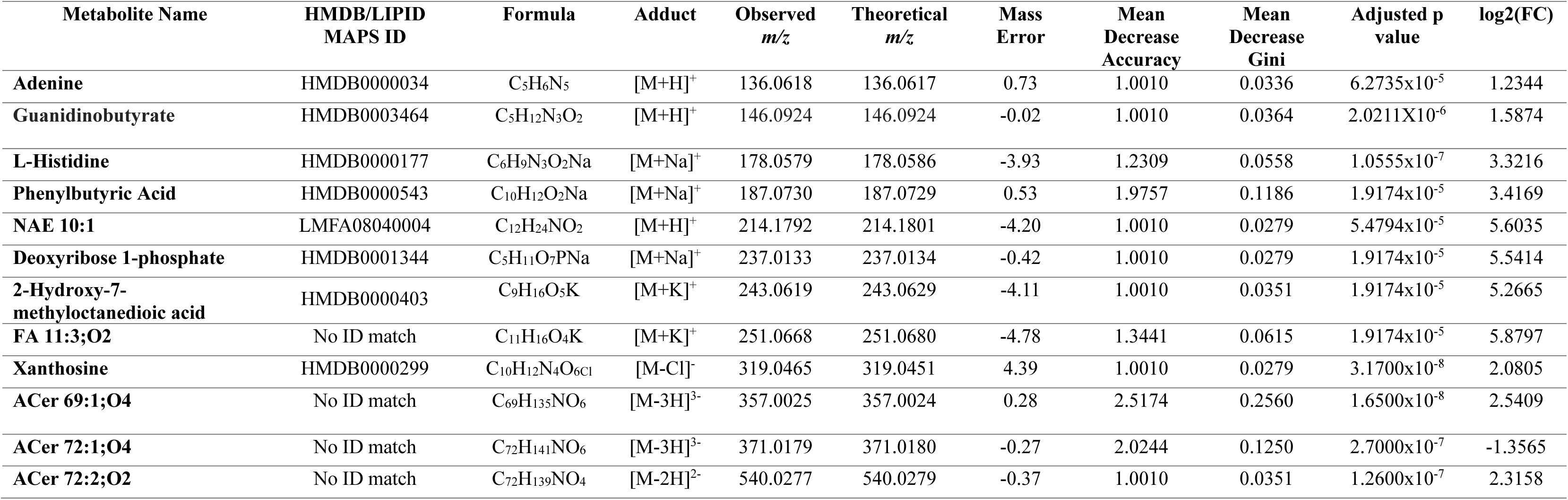
Significant metabolites identified in HA co-culture (*n*=10) with GIN cells (*n*=10). Metabolites were detected using AP-MALDI mass spectrometry with DHB matrix. Metabolites were selected using variable importance measures from random forest analysis (Mean Decrease Accuracy, Mean Decrease Gini), adjusted *p*-values from statistical testing, and log₂ fold change (log₂FC) values.

As shown in **Figure 8C**, we similarly examined the metabolic profiles of HA in co-culture relative to monoculture conditions. Astrocytes grown in monoculture clustered more tightly, consistent with a uniform metabolic phenotype in the absence of tumour-derived influence. In contrast, astrocytes in co-culture displayed broader variance and partial separation, particularly along PC2, suggesting metabolic change in response to the GBM microenvironment. Previous studies have shown that glioblastoma cells in co-culture with astrocytes can undergo transcriptional reprogramming, metabolic rewiring, and enhanced proliferative and invasive behaviours in response to astrocytic cues.^45^ However, delineating specific metabolic changes within each cell population remains experimentally challenging and is not yet well established. The partial overlap along PC1 indicates retention of core metabolic features, consistent with previous reports that astrocytes maintain fundamental phenotypic characteristics in glioma co-culture models reflecting shared metabolic profiles.^46^

To identify metabolites that were significantly altered in astrocytes during interaction with GBM cells, we compared metabolite intensities between HA cells in co-culture and monoculture (**Figure 8D**; **Table 1**). Several nucleoside and nucleotide-related metabolites, including xanthosine (*m/z* 319.0465), deoxyribose 1-phosphate (*m/z* 237.0133), and adenine (*m/z* 136.0618) showed significant increases in co-culture, implicating altered nucleotide turnover. Amino acid derivatives such as guanidinobutyrate (*m/z* 146.0924), histidine (*m/z* 178.0579), and phenylbutyric acid (*m/z* 187.073) were also elevated, potentially reflecting responses to oxidative stress, nitrogen metabolism, or protein turnover.^47,48^ Notably, a range of lipid-related metabolites including NAE 10:1 (*m/z* 214.1792), FA 11:3;O2 (*m/z* 251.0668), and long-chain ceramides such as ACer 69:1;O4 (*m/z* 357.0025), ACer 72:1;O4 (*m/z* 371.0179), and ACer 72:2;O2 (*m/z* 540.0277) were significantly more abundant in co-cultured astrocytes. These changes may suggest remodelling of membrane lipids and activation of signalling pathways linked to inflammation consistent with the established role of sphingolipid metabolism in GBM progression and evidence that astrocyte lipid metabolism contributes to brain pathology, including GBM.^49,50^

Comparison of the two cell populations in co-culture highlights potential metabolic crosstalk. Both GIN cells and HA cells in co-culture exhibited evidence of altered nucleotide turnover, with increased ribose-5-phosphate and glycerophosphoinositol in tumour cells paralleled by elevated xanthosine, deoxyribose 1-phosphate, and adenine in HA. Similarly, lipid-related changes were observed in both cells: GIN cells showed reduced ceramide accumulation, whereas astrocytes displayed increased abundance of NAE 10:1, unsaturated fatty acids, and long-chain ceramides, consistent with reciprocal remodelling of membrane and signalling lipids. Furthermore, detection of 4-hydroxybenzoic acid in GIN cells together with phenylbutyric acid enrichment in astrocytes points to complementary regulation of tyrosine metabolism across the two populations. This pattern aligns with reports of metabolic support from the tumour microenvironment, while the specific role of astrocytes in modulating amino acid availability remains to be determined.^51^

## Conclusions

This study establishes a microscopy-guided AP-MALDI-MSI workflow that enables single-cell metabolomic profiling while preserving cell-type identity within co-culture model. By integrating cell-specific labelling, high-resolution mass spectrometry, and a tailored annotation strategy, this platform allows untargeted, spatially resolved analysis of intercellular metabolic interactions.

Application of this workflow to glioblastoma–astrocyte co-cultures revealed distinct and complementary metabolic alterations across cell types, including altered nucleotide turnover, phospholipid and sphingolipid metabolism, and modulation of tryptophan and tyrosine pathways. These findings suggest coordinated metabolic exchange between glioblastoma and astrocytes within the brain microenvironment, potentially influencing tumour cell energy balance and providing new insight into how astrocytes contribute to tumour invasiveness. Integrating these findings with functional studies will further extend its potential to uncover the mechanistic underpinnings in disease progression.

Beyond the co-culture model, this workflow provides an accessible framework for investigating cell-cell metabolic interactions within more complex tissue environments and across diverse biological systems. Extending this approach to primary tumour samples and other multicellular models will allow direct exploration of metabolic relationships in their native microenvironments, advancing our understanding of how spatial metabolic organisation contributes to disease pathophysiology and therapeutic response.

## Associated Content

### Supporting Information

Supporting Information (PDF) includes **Table S1** listing biologically relevant metabolites identified by AP-MALDI using a DHB matrix and confirmed by LC-MS/MS based on retention time and fragmentation matching to reference standards, **Figure S1** showing the class distribution of LC-MS-identified metabolites, and **Figure S2** presenting statistical analyses of HA and GIN single-cell datasets.

### Author Contributions

The manuscript was written through contributions of all authors. All authors have contributed to the conception, data acquisition, and analysis, and have approved the final version of the manuscript.

### Funding Sources

This work was funded by the Biotechnology and Biological Sciences Research Council (Doctoral Training Programme) [grant number: BB/T008369/1] and a philanthropic donation from Sam White Legacy.

## Acknowledgments

The authors thank the technical staff at the Centre for Analytical Biosciences, University of Nottingham, for their support throughout this study. We also acknowledge Gilles Frache and Sue Kennerley from MassTech for their instrumental and technical assistance with the AP-MALDI-MSI system. We further acknowledge Dr. Roba Sofi (University of Bristol) for the development of the eGFP-expressing GIN glioblastoma cell lines used in this study.

## Data Availability Statement

The data that support the findings of this study are available from the corresponding author upon reasonable request.

## Table of Contents

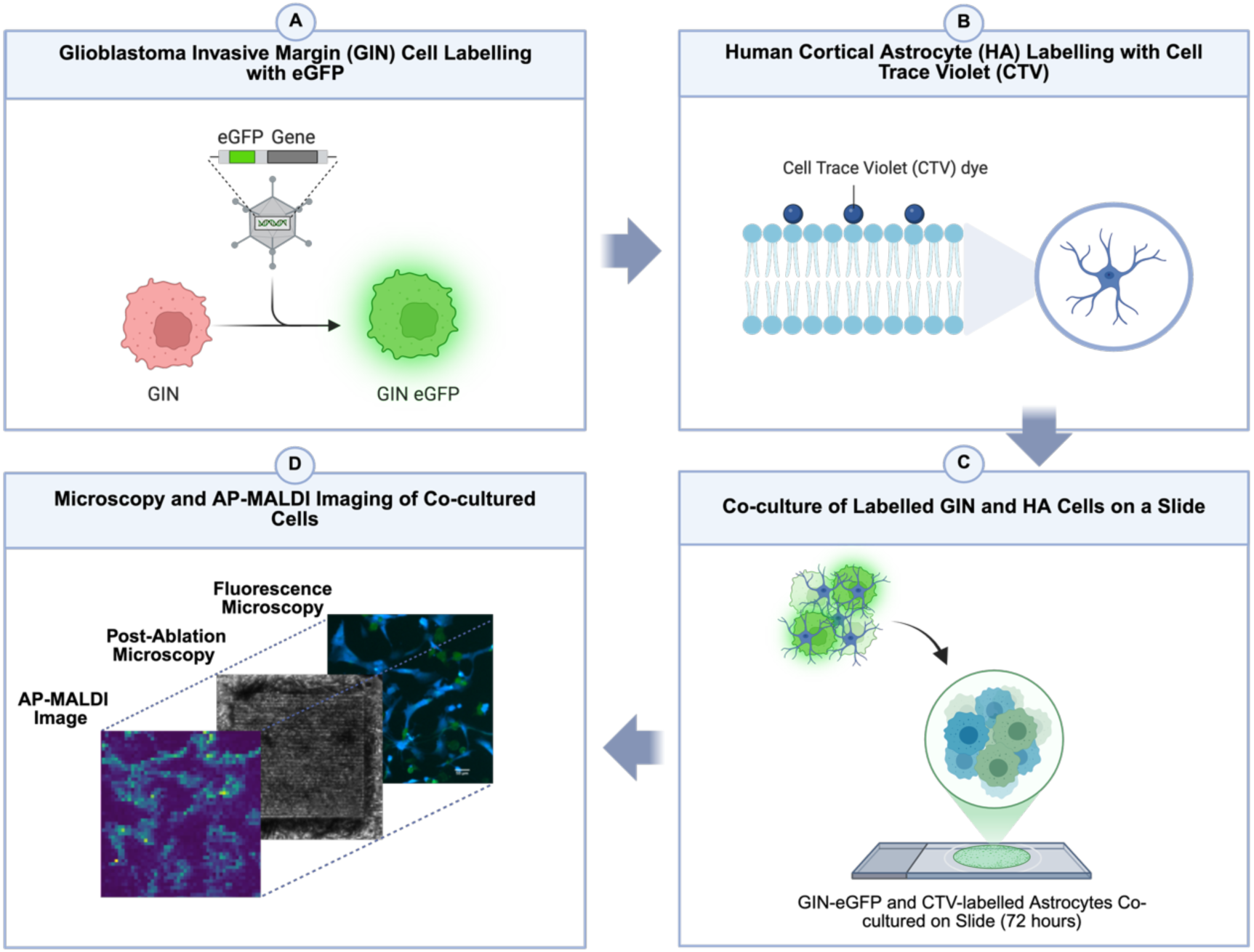

