## Supplementary document for "Single-Cell Metabolic Profiling in a Glioblastoma Co-culture Model Using AP-MALDI-based Mass Spectrometry Imaging"

| *Name* | *HMDB ID* | *Formula* | *Adduct* | *Observed m/z* | *Theoretical m/z* | *Mass Error* | *RT [min]* | *Level of Confidence* |
|---|---|---|---|---|---|---|---|---|
| ***Pyridoxine*** | HMDB0000239 | $C_8 H_{11} N O_3$ | $[M+H]^+$ | 170.0811 | 170.08173 | -0.63 | 7.48 | Level 1 |
| ***P-DMEA*** | HMDB0060244 | $C_4 H_{12} N O_4 P$ | $[M-H]^-$ | 168.0434 | 168.04167 | 1.73 | 12.55 | Level 4 |
| ***Nicotinamide*** | HMDB0001406 | $C_6 H_6 N_2 O$ | $[M+H]^+$ | 123.0552 | 123.0555 | -0.30 | 7.33 | Level 4 |
| ***N-Acetylputrescine*** | HMDB0002064 | $C_6 H_{14} N_2 O$ | $[M+H]^+$ | 131.1179 | 131.11815 | -0.25 | 16.80 | Level 2 |
| ***N-Acetyl-L-phenylalanine*** | HMDB0000512 | $C_{11} H_{13} N O_3$ | $[M-H]^-$ | 206.0826 | 206.0809 | 1.70 | 7.35 | Level 4 |
| ***N6-Acetyl-L-lysine*** | HMDB0000206 | $C_8 H_{16} N_2 O_3$ | $[M+ACN+H]^+$ | 189.1233 | 189.12352 | -0.22 | 11.21 | Level 4 |
| ***N-3-[(4-Acetamidobutyl)amino]propyl acetamide*** | HMDB0041947 | $C_{11} H_{23} N_3 O_2$ | $[M+H]^+$ | 230.1863 | 230.18654 | -0.24 | 15.03 | Level 4 |
| ***L-Valine*** | HMDB0000883 | $C_5 H_{11} N O_2$ | $[M-H]^-$ | 116.0720 | 116.07204 | -0.04 | 11.43 | Level 1 |
| ***L-Tyrosine*** | HMDB0000158 | $C_9 H_{11} N O_3$ | $[M-H]^-$ | 180.0669 | 180.06599 | 0.91 | 12.06 | Level 1 |
| ***L-Phenylalanine*** | HMDB0000159 | $C_9 H_{11} N O_2$ | $[M-H]^-$ | 164.0719 | 164.07131 | 0.59 | 9.92 | Level 2 |
| ***L-Norleucine*** | HMDB0001645 | $C_6 H_{13} N O_2$ | $[M+H]^+$ | 132.1019 | 132.10143 | 0.47 | 10.65 | Level 2 |
| ***L-Histidine*** | HMDB0000177 | $C_6 H_9 N_3 O_2$ | $[M+H]^+$ | 156.0768 | 156.07634 | 0.46 | 13.40 | Level 2 |
| ***L-Glutamine*** | HMDB0000641 | $C_5 H_{10} N_2 O_3$ | $[M+H]^+$ | 147.0764 | 147.07654 | -0.14 | 15.96 | Level 3 |
| ***Hypoxanthine*** | HMDB0000157 | $C_5 H_4 N_4 O$ | $[M+H]^+$ | 137.0458 | 137.0454 | 0.40 | 10.12 | Level 1 |
| ***Guanine*** | HMDB0000132 | $C_5 H_5 N_5 O$ | $[M+NH_4]^+$ | 152.0567 | 152.05668 | 0.02 | 11.83 | Level 4 |
| ***Cytosine*** | HMDB0000630 | $C_4 H_5 N_3 O$ | $[M+H]^+$ | 112.0505 | 112.05056 | -0.06 | 10.25 | Level 3 |

| | | | | | | | | |
|---|---|---|---|---|---|---|---|---|
| ***Cytidine*** | HMDB0000089 | $C_9 H_{13} N_3 O_5$ | $[M-H]^-$ | 242.0802 | 242.07217 | 8.03 | 12.31 | Level 4 |
| ***Creatinine*** | HMDB0000562 | $C_4 H_7 N_3 O$ | $[M+H]^+$ | 114.0661 | 114.06673 | -0.63 | 11.03 | Level 4 |
| ***Creatine*** | HMDB0000064 | $C_4 H_9 N_3 O_2$ | $[M+H]^+$ | 132.0767 | 132.07717 | -0.47 | 12.82 | Level 1 |
| ***Choline*** | HMDB0000097 | $C_5 H_{13} N O$ | $[M+H]^+$ | 104.1069 | 104.10775 | -0.85 | 19.33 | Level 2 |
| ***b-Ala-Lys*** | HMDB0060442 | $C_9 H_{19} N_3 O_3$ | $[M+H]^+$ | 218.1499 | 218.14991 | -0.01 | 13.45 | Level 4 |
| ***Adenine*** | HMDB0000034 | $C_5 H_5 N_5$ | $[M-H]^-$ | 134.0475 | 134.04549 | 2.01 | 8.90 | Level 4 |
| ***4-Hydroxybenzoic acid*** | HMDB0000500 | $C_7 H_6 O_3$ | $[M-H]^-$ | 137.0247 | 137.02273 | 1.97 | 4.78 | Level 4 |
| ***2-Furoic acid*** | HMDB0000617 | $C_5 H_4 O_3$ | $[M-H]^-$ | 111.0090 | 111.00683 | 2.17 | 14.76 | Level 4 |
| ***11-Nitro-1-undecene*** | HMDB0062669 | $C_{11} H_{21} N O_2$ | $[M+H]^+$ | 200.1644 | 200.16474 | -0.34 | 7.37 | Level 4 |
| ***2-Keto-glutaramic acid*** | HMDB0001552 | $C_5 H_7 N O_4$ | $[M-H]^-$ | 144.0305 | 145.0375 | 1.84 | 9.98 | Level 4 |

**Table S1.** List of biological metabolites identified using AP-MALDI with DHB matrix. Identification was confirmed by LC-MS/MS using retention time (RT) and fragmentation matching to reference standards.

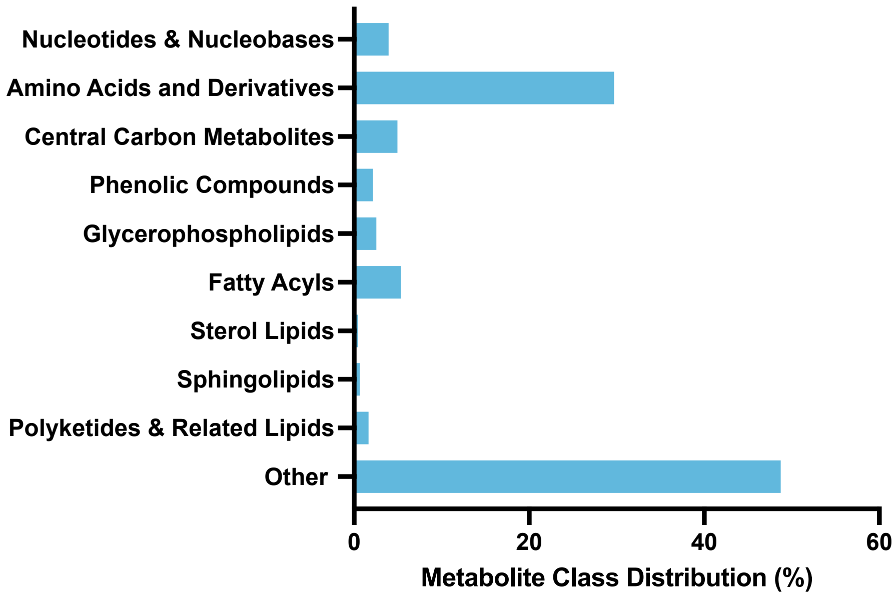


**Figure S1. Class distribution of LC-MS-identified metabolites.** Bar chart showing the percentage distribution of identified metabolites across major chemical classes detected by LC-MS analysis of $5x10^5$ cells. Classes include Nucleotides & Nucleobases, Amino Acids and Derivatives, Central Carbon Metabolites, Phenolic Compounds, Glycerophospholipids, Fatty Acyls, Sterol Lipids, Sphingolipids, and Polyketides & Related Lipids. The "Other" category contains metabolites that could not be confidently assigned to a specific class due to structural diversity or limited annotation. Percentages were calculated as the proportion of each class relative to the total number of LC-MS-identified metabolites.

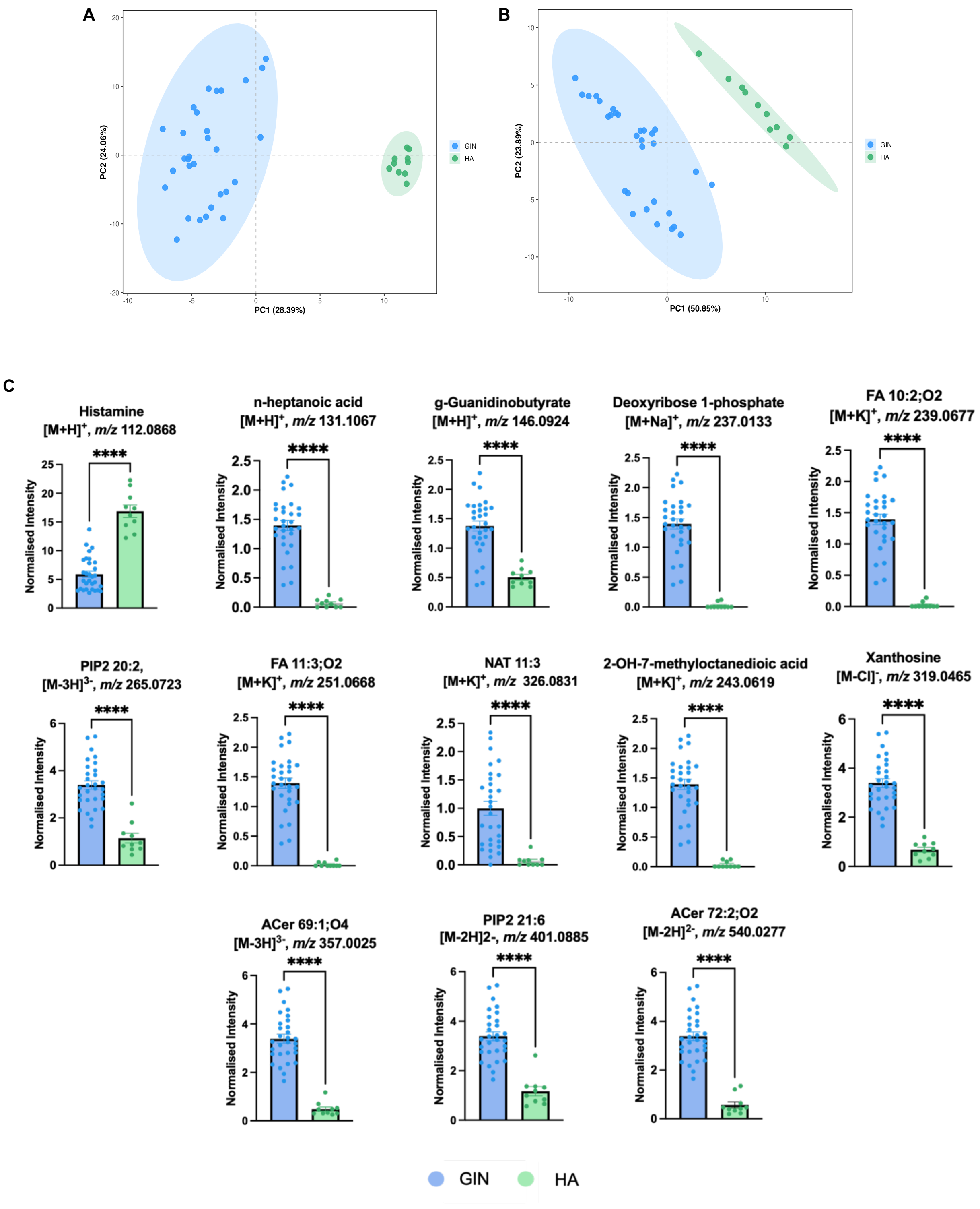


**Figure S2.** Comparative metabolomic analysis of HA and GIN single cells. PCA plot of HA and GIN single cells acquired in **(A)** positive ion mode and **(B)** negative ion mode. **(C)** Statistical comparison of metabolite profiles between GIN and HA cells. Significantly different metabolites were identified using univariate statistical analysis ($p < 0.05$) and Random Forest classification to assess feature importance.
